# SSA.ME Detection of cancer mutual exclusivity patterns by small subnetwork analysis

**DOI:** 10.1101/034124

**Authors:** Sergio Pulido-Tamayo, Bram Weytjens, Dries De Maeyer, Kathleen Marchal

## Abstract

Because of its clonal evolution a tumor rarely contains multiple genomic alterations in the same pathway, as disrupting the pathway by one gene often is sufficient to confer the complete fitness advantage. As a result mutated genes display patterns of mutual exclusivity across tumors. The identification of such patterns have been exploited to detect cancer drivers. The complex problem of searching for mutual exclusivity across individuals has previously been solved by filtering the input data upfront, analyzing only genes mutated in numerous samples. These stringent filtering criteria come at the expense of missing rarely mutated driver genes. To overcome this problem, we present SSA.ME, a network-based method to detect mutually exclusive genes across tumors that does not depend on stringent filtering. Analyzing the TCGA breast cancer dataset illustrates the added value of SSA.ME: despite not using mutational frequency based-prefiltering, well-known recurrently mutated drivers could still be highly prioritized. In addition, we prioritized several genes that displayed mutual exclusivity and pathway connectivity with well-known drivers, but that were rarely mutated. We expect the proposed framework to be applicable to other complex biological problems because of its capability to process large datasets in polynomial time and its intuitive implementation.

## Introduction

Because of internationally coordinated efforts such as TCGA^1,2^ and ICGC^3^, a vast number of cancer datasets are publicly available. Using these datasets to identify mutations and pathways driving cancer phenotypes has become an active field of research^4–7^. Tumorigenesis and tumor progression follow a clonal evolutionary model^8–11^. In this view, the disruption of a single gene in a molecular pathway often yields the complete fitness advantage associated with disruption of that pathway, making additional mutations in the same pathway redundant^8^. This evolutionary property can be exploited to understand cancer mechanisms by searching for patterns of genes that display mutual exclusivity (*i.e*. groups of genes which mostly have maximum one mutation per tumor). The identification of groups of genes showing patterns of mutual exclusivity across patients in large datasets has already been proven useful for the detection of driver mutations/pathways in single cancer types such as triple-negative breast cancer^12^, Lung Adenocarcinoma^13^ and in a pan-cancer setting^14,15^.

In practice the mutual exclusivity patterns are not always strict (hard patterns), *i.e*. most patterns occasionally show the presence of multiple mutations in a single tumor. This is possible because for example tumorigenesis can start in an initially less potent driver, but more potent drivers in the same pathway can accumulate at later times, providing an additional marginal beneficial effect (diminishing returns)^16^. Therefore, exploiting clonal behavior for identifying driver pathways requires searching for “soft” mutual exclusivity where two otherwise independent mutational events co-occur less than expected by chance^17^.

In order to discover genes that exhibit a mutual exclusive pattern in cancer, all possible sets of genes have to be examined. Due to the factorial computational complexity of this problem i.e. adding an extra gene to the pattern implies that the algorithm’s processing time increases factorially^18^, this problem cannot be solved for large data sets. Current methods mainly cope with this by prioritizing potential important genes upfront, filtering out genes which seem to be less important mainly based on the frequency with which they are mutated across tumor samples. Methods such as Dendrix^6^, CoMEt^19^ and Multidendrix^13^ explicitly try to find the largest set of genes that exhibit a mutual exclusivity pattern after a filtering step, using an integer linear program or a Markov chain Monte Carlo approach while methods such as MEMo^5^ and Mutex^20^ rely on the use of the human interaction network to further constrain the search space by using the knowledge that mutually exclusive genes are likely to be located in the same molecular pathways.

MEMo relies on a human protein-protein interaction network to search for the largest set of genes that are closely related in the network and that exhibit mutual exclusivity, whereas Mutex uses a directed signaling network. Although using a network restricts the search space, searching for patterns of mutual exclusivity is still a difficult task. For these reasons, both MEMo and Mutex require a stringent filtering of the input (the input of Mutex is required to consist of less than 500 genes, MEMo is capable to analyze about 250 genes). As a result, potential drivers that are rarely mutated are likely to be missed.

Therefore we developed SSA.ME (Small Subnetwork Analysis with reinforced learning for detecting Mutual Exclusivity patterns), a computational tool that searches for genes that belong to common patterns of mutual exclusivity and that are closely connected on an interaction network to prioritize drivers. It uses a novel methodology named Small Subnetwork Analysis with reinforced learning (SSA) that divides a complex problem, *i.e*. finding the largest set of genes that exhibit a mutual exclusivity pattern, into many simpler ones by calculating measures for mutual exclusivity in many small subnetworks. By solving these simpler problems iteratively, each time biasing the search space based on results of previous iterations, SSA.ME can prioritize potential driver genes with linear algorithmic complexity. This, in principle, allows it to process large input datasets in short computational times and therefore, in contrast to previous approaches, requires little prior filtering.

To assess the performance of SSA.ME we re-analyzed the breast cancer dataset from the 2012 cancer genome atlas (TCGA)^2^ without filtering the genomic variants up front. Despite adding many more mutations in the input, we could prioritize well-known drivers that are found to be recurrently mutated in different tumors. However, in addition to prior findings we could prioritize several genes that displayed mutual exclusivity and pathway connectivity with well-known drivers, but that were rarely mutated in the different tumors and therefore were missed by other methods that search for mutual exclusivity.

## Results

### SSA.ME Implementation

To identify cancer drivers we develop SSA.ME (Small Subnetwork Analysis for mutual exclusivity), a method that searches for small subnetworks of the interaction network containing mutated genes that show a pattern of mutual exclusivity. SSA.ME reformulates the complex problem of finding the largest set of mutually exclusive genes into many independent and less complex sub-problems. SSA.ME scores many small subnetworks for their potential to contain genes that belong to a mutual exclusivity pattern, instead of explicitly searching for the largest set of mutual exclusive genes. Using these small subnetwork scores in a reinforced learning framework allows prioritizing individual genes that are likely to belong to a mutual exclusivity pattern, without ever having to explicitly evaluate the largest set of mutually exclusive genes.

The method is outlined in Figure 1. SSA.ME searches the local neighborhood around a set of predefined seed genes. In this case, the seed genes correspond to all genes mutated in at least one sample. In each iteration step of the algorithm, genes in the neighborhood of a seed gene are selected into a small subnetwork with a chance proportional to their gene scores (which are chosen to be uniformly distributed in the first iteration). These small subnetworks are subsequently scored based on the mutual exclusivity signal of the genes in each small subnetwork. Individual gene scores are updated proportional to the mutual exclusivity scores of the selected small subnetworks to which they belonged. Updating of the gene scores modifies the likelihood with which each gene will be selected in subsequent iteration steps. The iterative process continues until the method converges to a solution or a maximum number of iterations is reached. The output of SSA.ME consists of an interactive network together with supporting files compatible with Cytoscape^21^.

**Figure 1.**
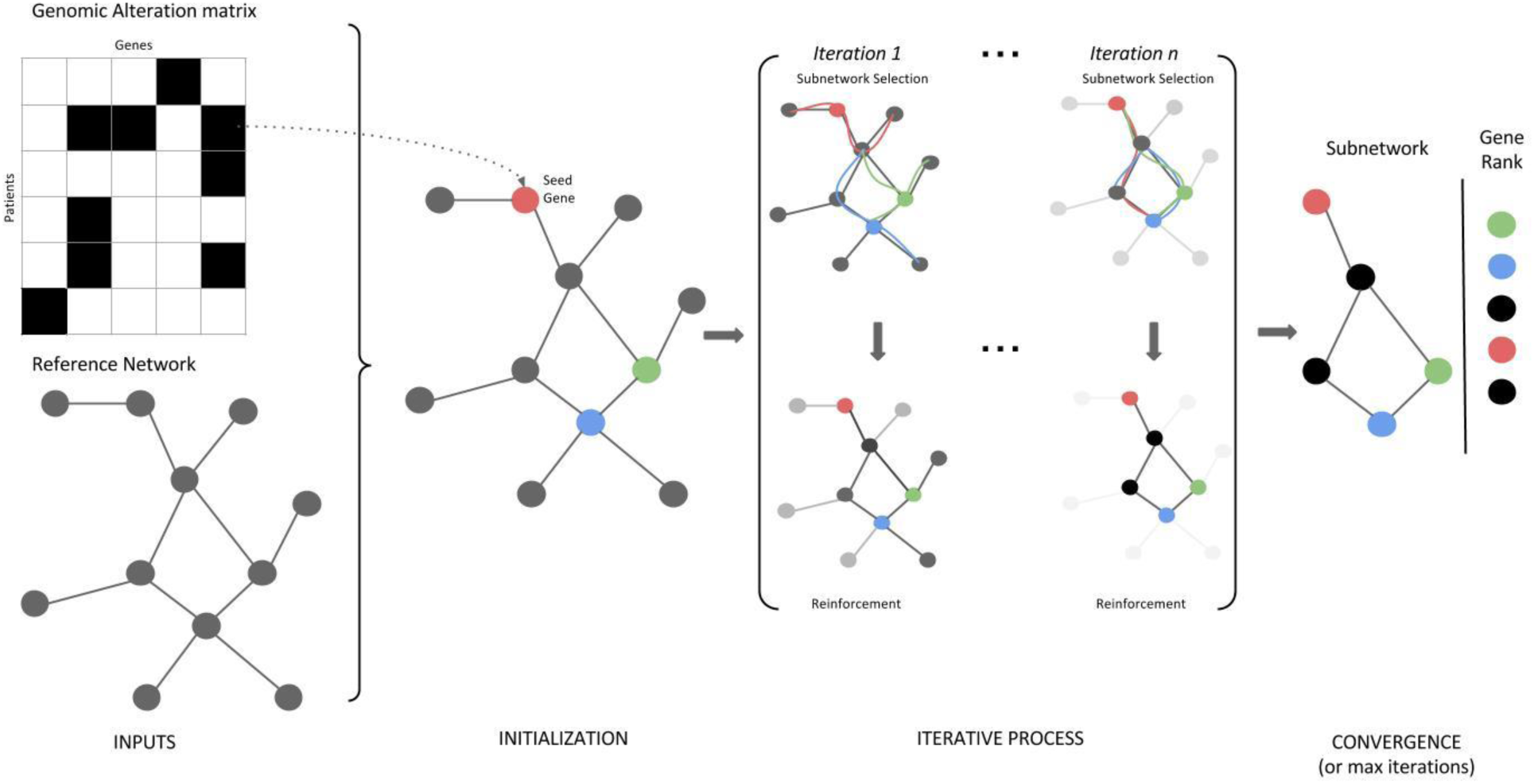
Overview of SSA.ME. The input consists of a matrix containing genomic alterations (i.e. mutations or copy number alterations, among others) across patients (depicted as black tiles) and a human reference network. In a first initialization step, every gene which has at least one genomic alteration across all patients is selected as a seed gene (colored genes in the network). The gene scores (represented as the opacity of the genes in the networks) are uniformly set to a value of 0.5. In every subsequent iteration step, small subnetworks will be generated, starting at every seed gene. Every gene adjacent to the small subnetwork has a chance proportional to its score to be incorporated in the small subnetwork. When a certain size has been reached the small subnetwork generation will stop and a score for each selected small subnetwork will be calculated based on the mutually exclusivity pattern found within this small subnetwork. At the end of every iteration step these small subnetwork scores will be used to update gene scores, altering the chance of genes to be incorporated into the small subnetwork in subsequent iteration steps. Upon convergence it can be seen that a few genes have high scores while others have scores close to 0. Genes are ranked based on their gene scores which reflects their potential to belong to a mutual exclusivity pattern.

## Performance on simulated data

To evaluate the robustness of the method with respect to the used reference network, we applied SSA.ME on a simulated dataset in combination with a high quality human reference network (see Materials and Methods) and underconnected/overconnected versions of this reference network (with respectively 10%, 25% and 50% of the network edges being deleted or added). Per network, 100 simulations were performed. Each simulated dataset contained a target mutual exclusivity pattern consisting of maximally 20 genes interacting on the reference network that were mutated in 30% of the samples (see Materials and Methods).

Applying SSA.ME on each simulated dataset resulted in a ranked gene list. The top x% of the gene list were considered as genes belonging to a mutual exclusivity pattern, whereas the remainder of the genes were considered not to exhibit mutual exclusivity. Performance was evaluated by plotting the sensitivity versus the specificity where the sensitivity is defined as the percentage of genes belonging to the target mutual exclusivity pattern that were retrieved amongst the x% highest ranked genes and the specificity, defined as the proportion of genes not present in the target pattern that were correctly classified as such. The results are shown in Fig. 2A for the highest ranked genes as this is the range that is of biological relevance (correctly identifying positives). The full ROC plot and the sensitivity/PPV plots can be found in the supplementary Fig. 1S.

**Figure 2.**
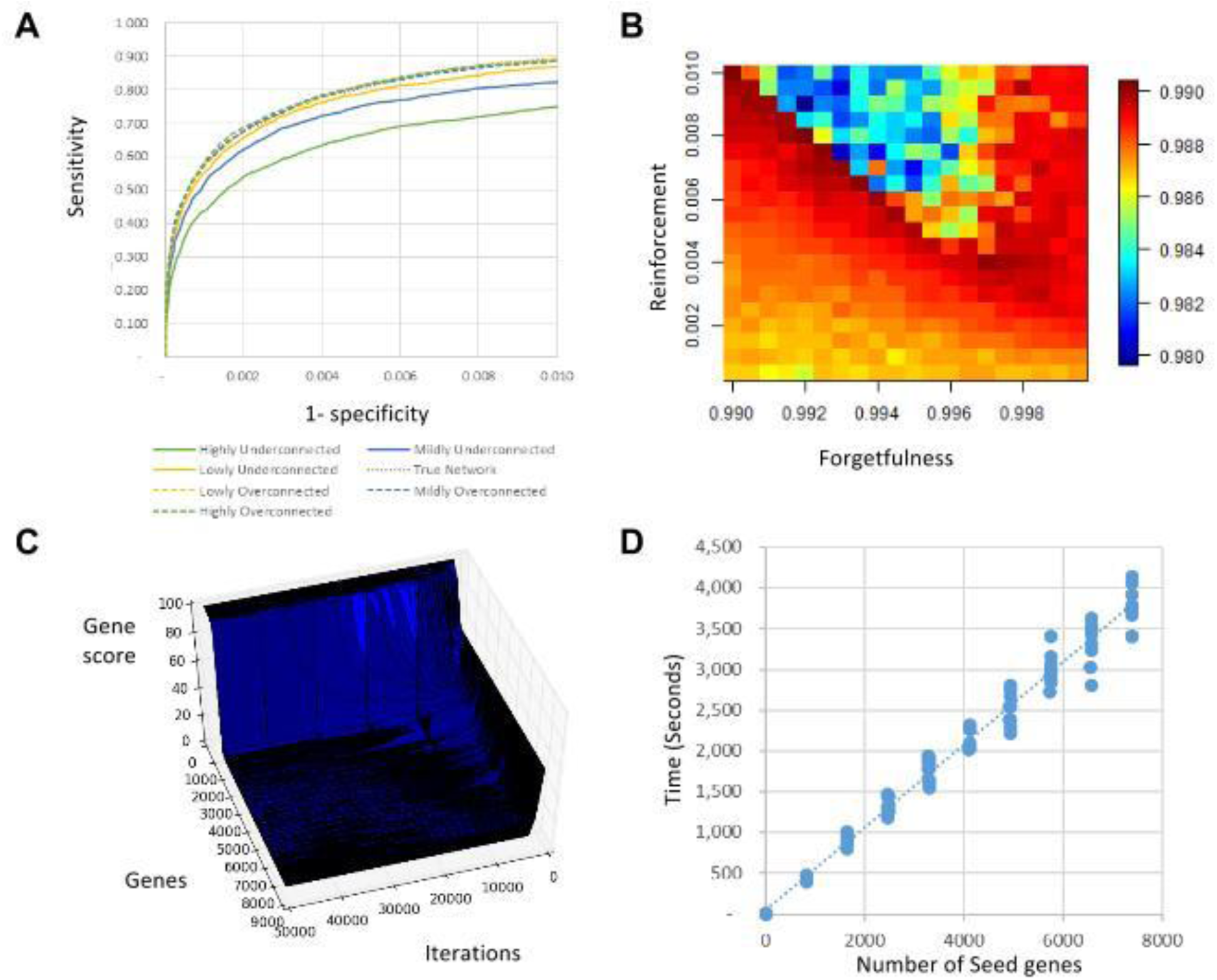
Performance on Simulated Data. **A)** Robustness of the predictions with respect to the used reference network. The X-axis represents 1-specificity and the Y-axis represents sensitivity. Underconnected networks lead to lower performance while overconnected networks result in similar, although lower, performance to when using the the true network. Note that, for clarity reasons, the range of the x-axis is restricted to [0, 0.01]. **B)** Heat map depicting parameter sensitivity. AUC values for every analyzed parameter pair are depicted. Warm colors depict higher AUC values while cold colors depict lower AUC values. It can be seen that the best performance is achieved on the diagonal for combinations of reinforcement and forgetfulness of 1. **C)** Plot visualizing convergence and stability of convergence of gene scores. The X-axis represents the number of performed iterations, the Y-axis displays all genes in the reference network (black lines in the plot) and the Z-axis represents the gene scores. All genes start on the right side with a gene score of 0.5. Most of them converge fastly to 0 or 1. As no inflecting lines are observed, convergence is stable. Results shown on a plot depicting scores for all genes at every iteration step. **D)** Plot showing linear time complexity of the algorithm with respect to the number of seed genes. Each dot on the plot represents the time to convergence of a separate run. Per tested number of seed genes, 10 simulations were performed. Results were obtained by running the algorithm on one single processor Intel(R) Xeon(R) CPU E5-2670 0 @ 2.60GHz.

Figure 2A indicates that the best performance is obtained using the reference network without added or deleted edges, as for the same relative increase in sensitivity less false positives are predicted (lower relative increase in 1-sensitivity). The method shows in general a high resilience of the results to using an overconnected network. In this case the method is capable of successfully prioritizing most of the mutually exclusive genes with a low number of false positives (which is the range we envisage when only showing the values of the 1-specificty between 0 and 0.01). With an underconnected network the maximal sensitivity that can be reached will get restricted as some of the genes that show mutual exclusivity can no longer be connected in the network.

To assess the sensitivity of the method versus its parameter settings we ran SSA.ME on the same simulated data each time using a different combination of the reinforcement and forgetfulness parameters. Hereby reinforcement values were varied from 0.0005 to 0.0100 in steps of 0.0005. Forgetfulness values varied from 0.99 to 0.9995 in steps of 0.0005. Note that values of the forgetfulness closer to 1 imply that less is ‘forgotten’ and values of reinforcement are consistently lower than the ones of the forgetfulness to ensure that only few true positives will be reinforced. For each parametercombination 10 simulated datasets were analyzed. The performance per parameter combination was assessed using the area under the ROC (Fig. 2B). In general a low performance is obtained if the forgetfulness is relatively low compared to the reinforcement. In those settings false positives might become reinforced relatively more than some weak or isolated true positives. However, in ranges where the forgetfulness is close to 1, the performance is more robust to the choice of the reinforcement value. Best performances were obtained on the diagonal where irrespective of their absolute values the sum of the values of *r* and *f* are close to each other *r + f* ≈ 1. In most cases, a combination where the sum of the reinforcement and the forgetfulness is higher than one results in lower performances because then again the reinforcement becomes relatively high compared to the forgetfulness, resulting in relatively more false positives.

To show that the method converges to a stable solution, we ran it on one simulated dataset for 50.000 iterations. Fig. 2C shows that the method exhibits a consistent behavior, i.e. after a gene obtains a high gene score, it will remain consistently high or vice versa. Furthermore this figure shows that the algorithm converges, provided a sufficient number of iterations have been performed.

To analyze its complexity with respect to the number of seed genes, we ran SSA.ME on 10 different simulated dataset, each time using an increasing number of seed genes (ranging from 1 to 8000 genes). Datasets contained incrementally more added seed genes. Seed genes were added gradually according to the frequency with which they were found mutated in the different tumor samples, hereby assuming that the most frequently mutated genes are the ones that in a real setting would also be prioritized as the most promising seeds. These runs were repeated on 10 different simulated datasets. Results are visualized in Fig. 2D and clearly show the linear complexity of the algorithm with respect to the number of seed genes.

### Analysis of the TCGA breast cancer data

To test the biological relevance of the predictions we applied SSA.ME on the well-studied TCGA Breast Cancer 2012 dataset^2^ using a high quality human reference network (see Materials and Methods). As seed genes we used all genes carrying somatic mutations or copy number alterations, provided the latter alterations also resulted in positively correlated expression values of those copy number altered genes. After running SSA.ME, genes were ranked according to their gene score and the highest ranked genes were prioritized as putative drivers. The cut-off on the ranked list was chosen so that, given a set of known cancer genes, a good trade-off between sensitivity and precision was obtained, i.e. we use the cut-off so that a maximal sensitivity was obtained with a PPV higher than 80% (Fig. 3A). Note that the PPV represents a lower boundary on the actual number of true positive predictions as all genes not previously associated with cancer according to CGC^22^, Malacard^23^ or NCG^24^ are regarded as false positives. Applying SSA.ME on the TCGA BRCA 2012 dataset resulted in 49 genes being prioritized as potential breast cancer associated genes. Because of the nature of the method this prioritized gene list contains both putative drivers, but also ‘linker genes’ that connect genes that are part of a mutual exclusivity pattern but that are not mutated themselves. These ‘linker genes’ are therefore no true drivers, but have driver potential as they were found in the network neighborhood of true drivers.

**Figure 3.**
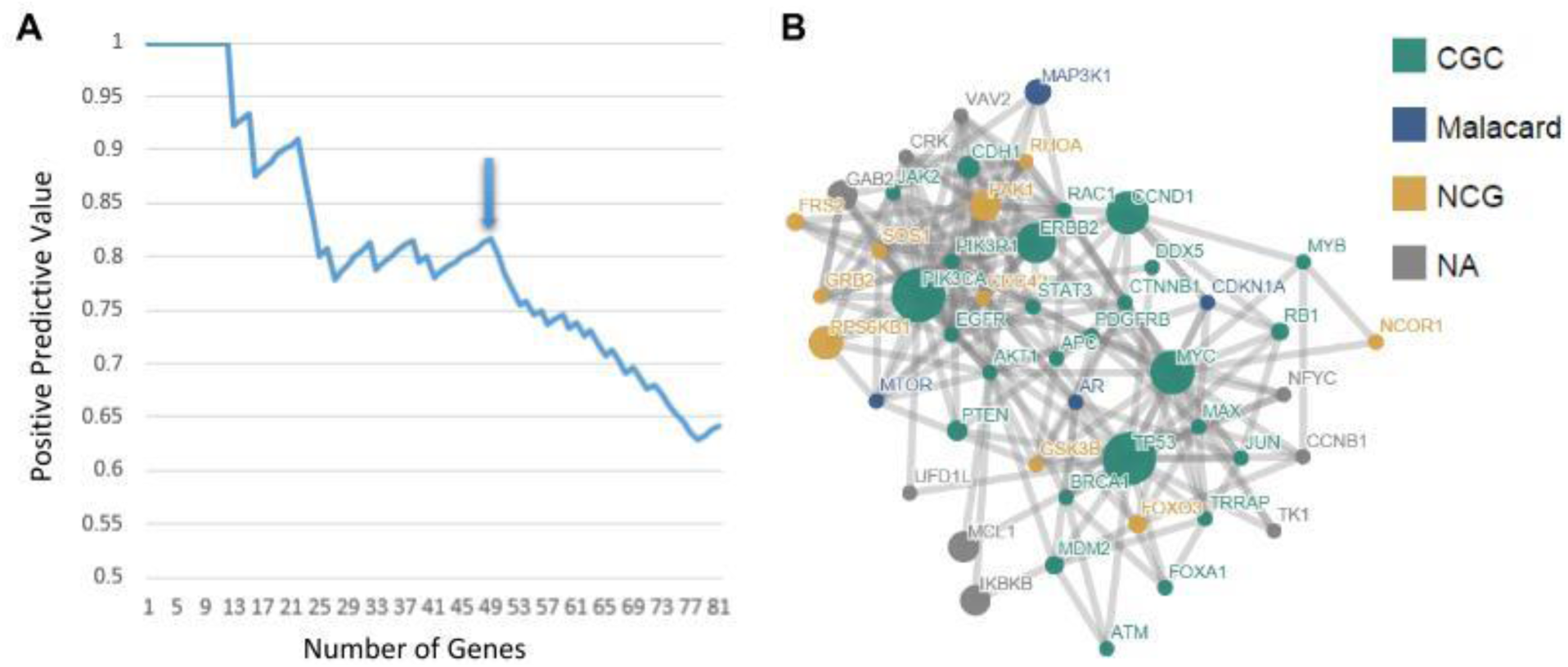
Application of SSA.ME on TGCA Breast Cancer dataset. **A)** Determination of the number of genes to be prioritized as cancer drivers. Genes were ranked according to their gene score obtained with SSA.ME. The X-axis represents the number of genes in the list of prioritized genes obtained by setting a cut-off on the rank. The Y-axis represents the positive predictive value (PPV) for the genes present in each list that corresponds to a given rank threshold. At the chosen threshold (arrow) 49 potential cancer drivers were prioritized. **B)** Subnetwork obtained after using SSA.ME on the TGCA breast cancer dataset. Seed genes and network were as defined in the main text. Genes are represented by nodes. If the gene had been associated with cancer, this is indicated by the color of the database in which the association was described. Gray genes correspond to genes not present in the Census of Cancer Genes, Malacard or the Network of Cancer Genes database. The size of the node reflects the number of samples in which a gene was found mutated.

The subnetwork in Fig. 3 displays the 49 prioritized genes and all edges in the reference network connecting them. Most of these genes (40 out of 49) have previously been associated with either breast cancer or cancer in general (Supplementary Table 1). 5 genes of those 40 were selected as ‘linker genes’ (i.e. they did not display alterations in the breast cancer dataset), but have been associated with other cancer types (i.e., *CDC42, CDKN1A, RAC1, GSK3B* and *CTNNB*). This indicates linker genes are potential cancer drivers.

Of all 49 predictions, 9 genes were not previously associated with breast cancer in Malacards^23^, or associated with cancer in general by the Cancer Gene Census^22^ or Network of Cancer Genes^24^ (*CCNB1, CRK, GAB2, IKBKB, MCL1, NFYC, TK1, VAV2* and *UFD1L*). Of those genes, two were selected as ‘linker genes’ (CRK and TK1). Of the remaining 7 genes, 4 were missed by previous analyses on the same TCGA breast cancer dataset because they were mutated in less than 2% of the samples and therefore did not pass the filtering strategies commonly applied prior to the driver analysis (i.e. *VAV2, UFD1L, NFYC*, and CCNB1). The genes IKBKB, MCL1 and GAB2 pass the filtering criteria used by previous methods. Of those IKBKB was also detected in the original TCGA analysis, reported to belong to a pattern of mutual exclusive genes based on the MEMo analysis, whereas MCL1 and GAB2 were not.

We could show that the mutations carried by the 49 prioritized genes and by the set of 9 cancer related genes not previously reported in cancer reference databases followed a CADD^25^ score distribution significantly different from the CADD score distribution of mutations in non-cancer related genes (Fig. 4A), pointing towards the functional relevance of at least some of the mutations carried by the predicted drivers. In addition, we could find clear associations for the novel predictions with cancer in literature (see Supplementary Note). Although at least visually for some of these driver candidates, the mutations they carry seem to cluster at the same genomic positions (Fig. 4B and Fig. 2S), none of them scores highly significant for clustering of their mutations according to the results provided in the pan-cancer analysis^1^ or the results we obtained by running SomInaClust^26^ (see Supplementary Note). This indicates that indeed without using network-based information, it would be difficult to prioritize these rarely mutated genes.

**Figure 4.**
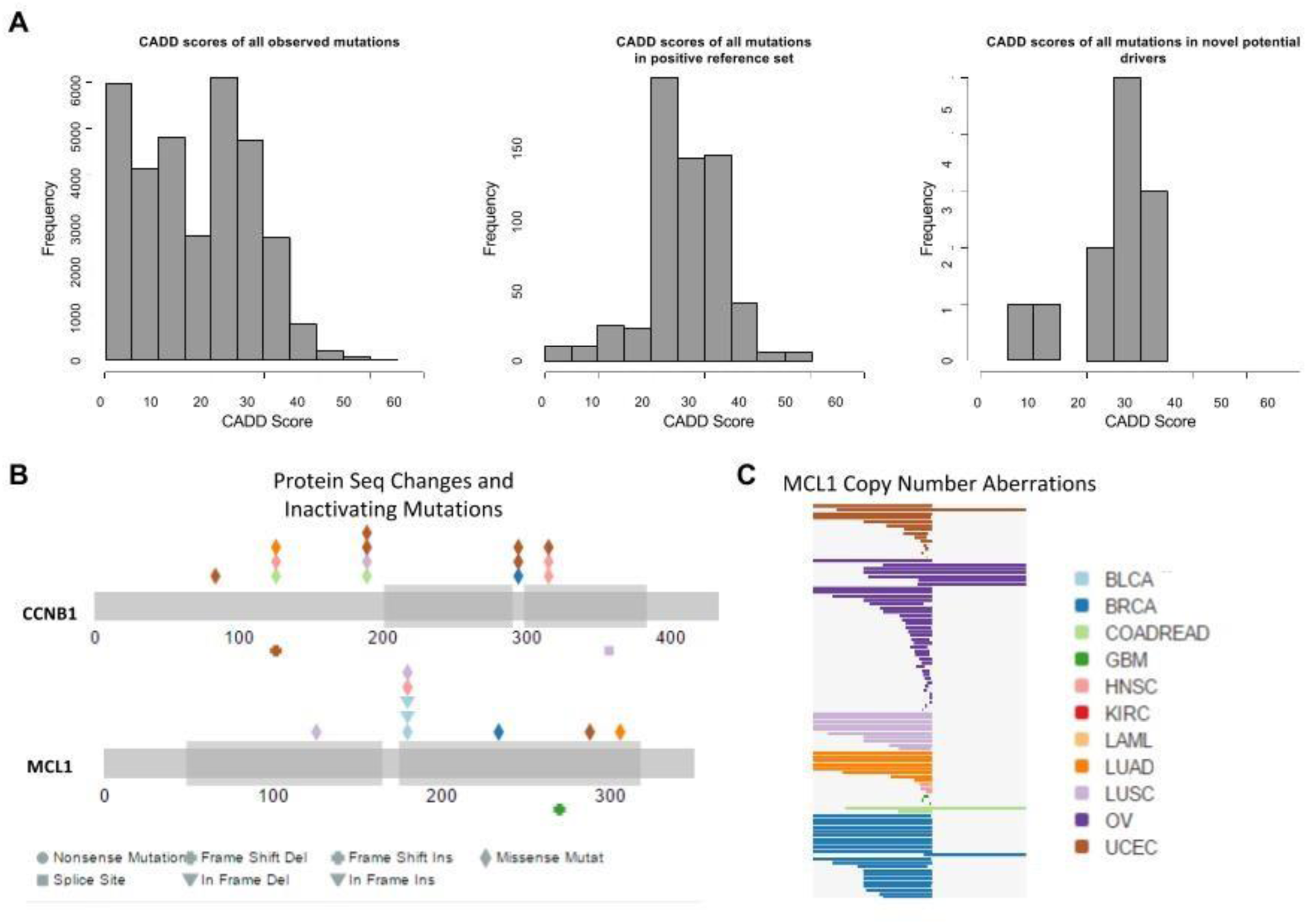
Analysis of selected genes. **A)** CADD score distribution of mutations of the unselected set (left histogram), the positive set (middle histogram) and the set containing the mutations in the novel predicted driver genes (right histogram). The X-axis depicts the CADD score and the Y-axis depicts the frequency of mutations having a CADD score within a certain range. **B)** Positioning of the mutations found in the TCGA Pan-Cancer dataset along the CCNB1 and MCL1 gene loci. Only a subset of copy number aberrations are included in this representation for MCL1. Figure obtained with MAGi. **C)** Copy number aberrations observed in the MCL1 gene in the TCGA Pan-Cancer dataset. Figure obtained with MAGi^35^.

## Comparison with TCGA Analysis

We compared the previously obtained predictions^2^ of MEMo, with our predictions. To maximize comparability between our results and those of MEMo on the same TCGA dataset, we reproduced the same filtering approach and network of the original breast cancer study and ran SSA.ME (see Materials and Methods).

Because of the high similarity of the mutual exclusivity patterns detected by MEMo in the original paper (patterns consisting of maximally 8 genes that varied in most cases in no more than one gene), we collapsed the 23 genes of all patterns and depicted them as a network (Fig. 5A). The subnetwork obtained by SSA.ME using the same filtered dataset consists of 33 genes (applying the same cut-off criteria as mentioned above) of which 18 were also found in the MEMo network (Fig. 5B). 5 genes retrieved by MEMo were not detected by SSA.ME (*NBN*, *CHECK2* and *MDM4*) as they were no longer present in the filtered list we used as input, whereas they must have been present in the original input of MEMo: in contrast to what has been described in the original TGCA paper we found these genes to be mutated in less than 2 samples and therefore removed them from our analysis. The score of *ATM* just fell below the chosen threshold (it ranked 36 with SSA.ME and the cut-off we used was set at 33) and was therefore also missing from our prioritization. *ATK3* was truly missed in our analysis as the small subnetworks to which it belonged never received consistently high scores during subsequent iteration steps.

**Figure 5.**
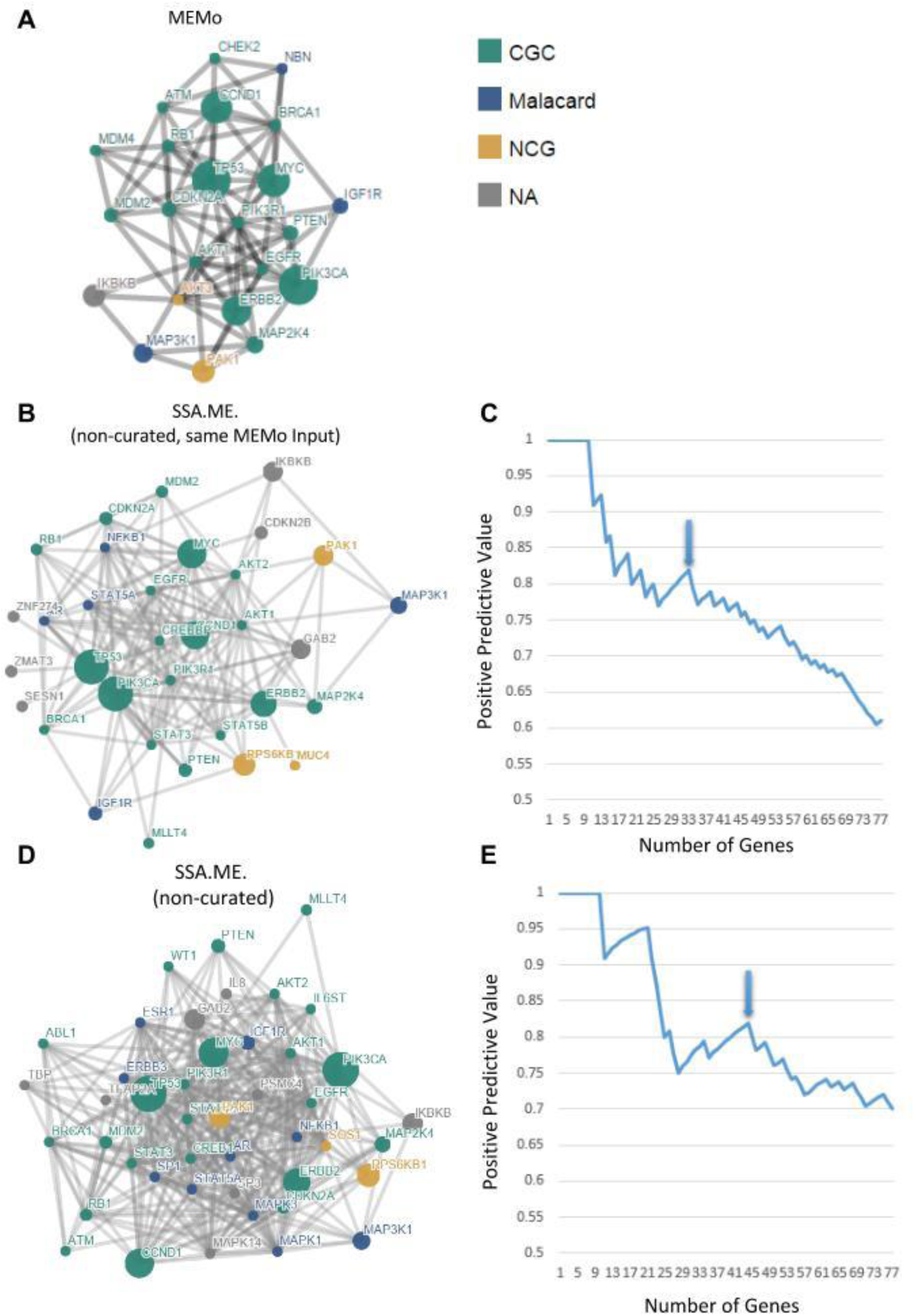
Comparison between SSA.ME and MEMo. Prioritized driver networks obtained by MEMO as retrieved from the original mutually exclusive modules outlined in the breast cancer TCGA paper (Panel **A**), obtained by SSA.ME using the filtered data (Panel **B**), and using the non-filtered data as input (Panel **D**). Genes are represented as nodes. Colors refer to the databases in which associations of the indicated genes with breast cancer or cancer have been described. Gray genes were not found to be associated with breast cancer/cancer according to the used reference databases. Panel **C** and **E** represent the PPV analysis of results obtained by applying SSA.ME on respectively the filtered and non-filtered datasets. Y-axis represents the PPV according to the reference databases. X-axis represents the number of genes in lists of prioritized genes of increasing order. Sizes of gene lists were determined by ranking genes according to their gene scores and counting the number of genes with a rank lower than a given threshold. Arrows indicate the thresholds that were chosen to select the genes in the represented networks. We choose the threshold on the ranked list so that a good trade-off between sensitivity and precision was obtained given a set of known cancer genes, i.e. we use the cut-off so that a maximal sensitivity was obtained with a PPV higher than 80%.

On the other hand we found 10 additional genes that were not retrieved by MEMo (of these, 4 were also found using the high quality network in the analysis described above: *AR, STAT3, RPS6KB1* and*GAB2*). Some of the additional genes had previously been associated to breast cancer (*AR* and *ESR1*) or to cancer in general (*MUC4* and *CCDN1*). The reason why we detect more genes than MEMo is partially due to the choice on the cut-off, but also because of the inherent differences in selection criteria between the methods: first, our method only requires that the selected genes are members of the local neighborhood of genes that exhibit a mutual exclusivity pattern across patients. Second, our method does not require stringent filtering which leaves the possibility of selecting rarely mutated genes.

These results thus show that SSA.ME is able to reproduce largely the same results as MEMo, provided the same input data are used or said otherwise genes that are highly ranked by MEMo are also highly ranked by our method. The discrepancies between the number of driver genes detected in this comparative analysis and the one above are due to the differences in the used networks. Above we choose to use a higher quality human network to reduce the possibility of including false positive interactions, whereas here we used the more connected interaction network used in the original TCGA dataset.

As shown above, the advantage of SSA.ME is that because of its reduced computational complexity it does not require stringent prior filtering of the data and therefore can also predict cancer drivers that are, for instance, infrequently mutated across the different samples. One could argue that not filtering the data can deteriorate the results as having more potentially false positives in the input list could dilute the true signals in the data and prevent the method from finding these true positives. To prove this is not happening we also applied SSA.ME on the less filtered data using the same reference network as in the original TCGA paper^2^. Less filtered data here correspond to all genes having at least one genomic alteration (5641 genes). Applying SSA.ME to these data and applying the same criteria to set the cut-off as mentioned above resulted in a driver network of 44 genes being selected from a total 5641 of genes (Fig. 5D). For comparison, with the filtered input set, 33 genes were prioritized out of the 237 using the same heuristics to set the cut-off on the size of the prioritized gene lists. Assuming that filtering already prioritizes the most frequently mutated genes and thus the most promising candidates, we can argue that the list obtained with the filtered input is the most reliable. Remarkably, of the 33 genes prioritized with the filtered input, 27 also occurred amongst the 44 genes prioritized with the unfiltered set. This indicates that despite the much larger number of input genes, the presumably true signals in the data are still best recovered and, compared to the much larger input that was used only relatively few additional candidates are prioritized. Not relying on pre-filtering in contrast offers the additional advantage of also recovering candidates that would not have passed the standard used stringent filtering criteria.

## Discussion

We introduce SSA, a small subnetwork analysis technique with reinforced learning which solves a complex combinatorial search over an interaction network by calculating measures for mutual exclusivity in many small subnetworks of the interaction network. The method can be generically applied to any problem in which local neighborhoods in a network hold useful information.

Here we applied SSA to prioritize cancer driver genes that are located in each other’s neighborhood on the interaction network and of which the genetic alterations display patterns of mutual exclusivity across different tumor samples (referred to as SSA.ME). To overcome the inherent high algorithmic complexity posed by its combinatorial nature, the problem of identifying drivers is iteratively solved and in each iteration multiple small subnetworks are independently analyzed. All results of these small subnetwork analyses are used in subsequent steps to bias the search space. The advantage of splitting the complex problem into multiple less complex problems, is that SSA.ME is not restricted by the number of mutated genes in the input data. As such by circumventing the stringent filtering strategy that is required by most other methods to enable the search for mutual exclusivity, SSA.ME can identify drivers carrying rare mutations and is able to identify genes based on relatively small-sized tumor cohorts of which the genetic variants cannot be pre-filtered based on the mutation frequency across samples.

Because we never explicitly evaluate the largest set of mutual exclusive genes in the interaction network, the prioritized mutated genes are not guaranteed to be all mutually exclusive or to all belong the same network neighborhood. SSA.ME rather prioritizes genes that belong to local mutual exclusivity patterns. If one would be interested in finding the largest set of mutual exclusive genes or independent modules, the prioritized gene list would suffice as input for *de novo* discovery methods for mutual exclusivity, such as Dendrix, Multidendrix or CoMEt. However, given the incompleteness of the interaction network and the biology of clonal systems, imposing too strong global constrains, e.g. requiring that all genes belonging to a mutual exclusivity pattern should also all be closely connected in the network, might reduce the number of selected potential driver genes. This because patterns of mutual exclusivity can be broken because genes belonging to a specific pattern can be unconnected in the interaction network due to missing interactomics data. In addition, if mutations trigger different adaptive pathways that are, when occurring in the same tumor, synthetically lethal, the genes carrying the mutually exclusive mutations would belong to different local regions in the interaction network (incompatible pathways) that cannot co-occur in the same tumor.

We showed that the results obtained by SSA.ME were largely consistent with those obtained by MEMo on the same TCGA 2012 dataset^2^ and that the use of a stringently versus an non-stringently filtered input set did not deteriorate the quality of those findings. By applying on the same breast cancer dataset SSA.ME with mutational data that were not a priori filtered based on recurrence across samples, we could show the potential of the method in discovering rarely mutated driver genes. In addition to drivers reported in well-known databases, our method prioritized an additional 9 drivers, not yet covered by conventional cancer databases. Several of these additional drivers were found to be infrequently mutated in the breast cancer and pan-cancer datasets and therefore have been missed by statistical prioritization methods that rely on recurrence of mutations across different tumors.

Conclusively, SSA.ME allows exploiting network connectivity and mutual exclusivity to identify drivers. Because of its computational efficiency, it can be used without relying on mutational recurrency based information and as such allows for the detection of infrequently mutated drivers.

## Methods

### SSA.ME

Small Subnetwork Analysis with reinforced learning for detecting Mutual Exclusivity patterns (SSA.ME) is an algorithm that uses a reference network to detect mutual exclusive gene patterns in cancer. To accomplish this, SSA.ME performs two independent functions in an iterative manner: small subnetwork selection/scoring and reinforced learning. Each gene (node) in the reference network is initialized with an initial uniform gene score. Then, iteratively: starting from a set of seed genes, small subnetworks are selected favoring genes with high gene scores. Each selected small subnetwork is then scored based on how well the genes composing the small subnetwork belong to a mutual exclusivity pattern. Genes that consistently belong to small subnetworks with high scores thus exhibit mutual exclusivity with other genes in their neighborhood very well and are more likely to be selected in subsequent iterations. This will lead to high gene scores for genes which are part of a mutual exclusivity pattern. The pseudocode describing the algorithm can be found in Fig. 6.

**Figure 6.**
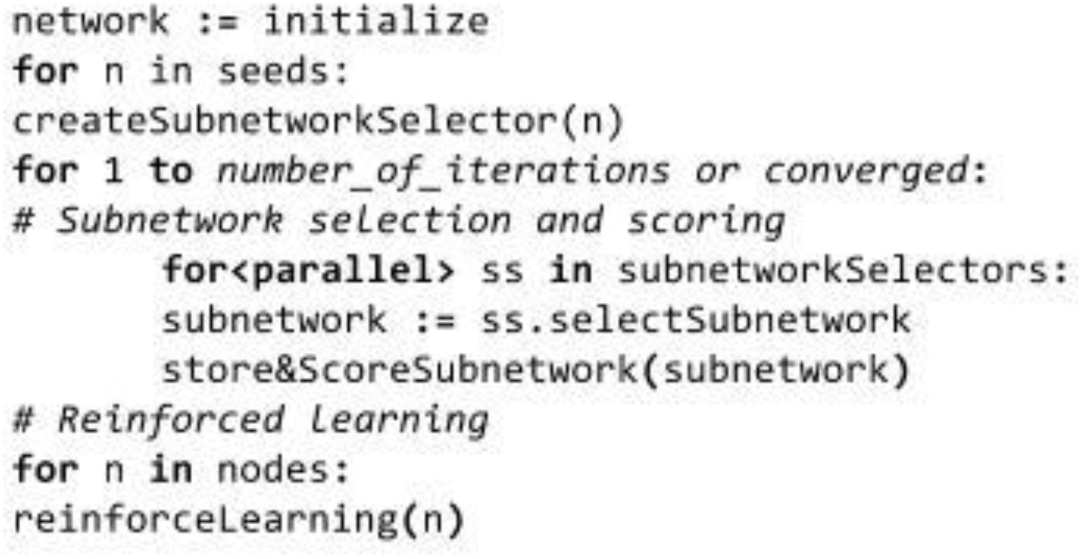
Pseudocode of SSA.ME algorithm

#### Initialization

The algorithm is initialized by giving each gene (node) a uniform initial gene score of 0.5. A static list of seed genes is defined that contains genes that possibly belong to a mutually exclusive pattern. Any type of biologically relevant filtering can be used to generate such gene list. In the context of this paper, seed genes are defined as all genes that were found to be mutated in at least one sample (tumor).

#### Small subnetwork selection and scoring

In each iteration small subnetworks of equal size are selected. Starting from every seed gene, subnetworks are selected by subsequently adding a gene adjacent to the current subnetwork. In order to be able to detect mutual exclusivity patterns of different sizes, the size of the small subnetworks varies from 3 to 6 genes between iterations. The probability of adding an adjacent gene to a small subnetwork is proportional to the gene scores of adjacent genes, expressing the assumption that mutually exclusive genes are likely to be located in the same adaptive pathway. Once constructed, each small subnetwork receives a mutual exclusivity score (MES). Each sample contributes to this score with a weight that is inversely related to the number of genes from the small subnetwork that were found mutated in that sample. This is calculated using the following equation:

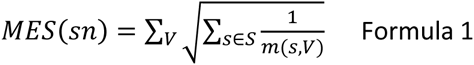

Where *V* are the genes present in small subnetwork *sn* ordered according to the number of samples in which these genes were found to be mutated. *S* is the set of samples pending to contribute to the mutual exclusivity score. Initially *S* includes every sample with a mutation in one of the genes in the small subnetwork, but every time a sample is used to calculate a mutual exclusivity score it is removed from *S*. In this way a sample can only contribute once to the *MES. m*(*s, V*) is the number of genes in *V* which are mutated in sample *s*. This value would be equal to 1 if the genes in gene set *V* are all members of a perfect mutual exclusive pattern and |*V*| if all genes in *V* are mutated in all samples. The square root allows giving relatively higher mutual exclusivity scores to small subnetworks for which each gene is mutated in approximately the same number of samples.

Next, the *MES* are ranked from highest to lowest and their ranks are divided by the maximum rank (Fig. 7). We end up with a ranked *MES* (*rMES*) between zero and one where zero refers to the small subnetwork having the least evidence for mutual exclusivity and one refers to the small subnetwork having the most evidence for mutual exclusivity.

**Figure 7.**
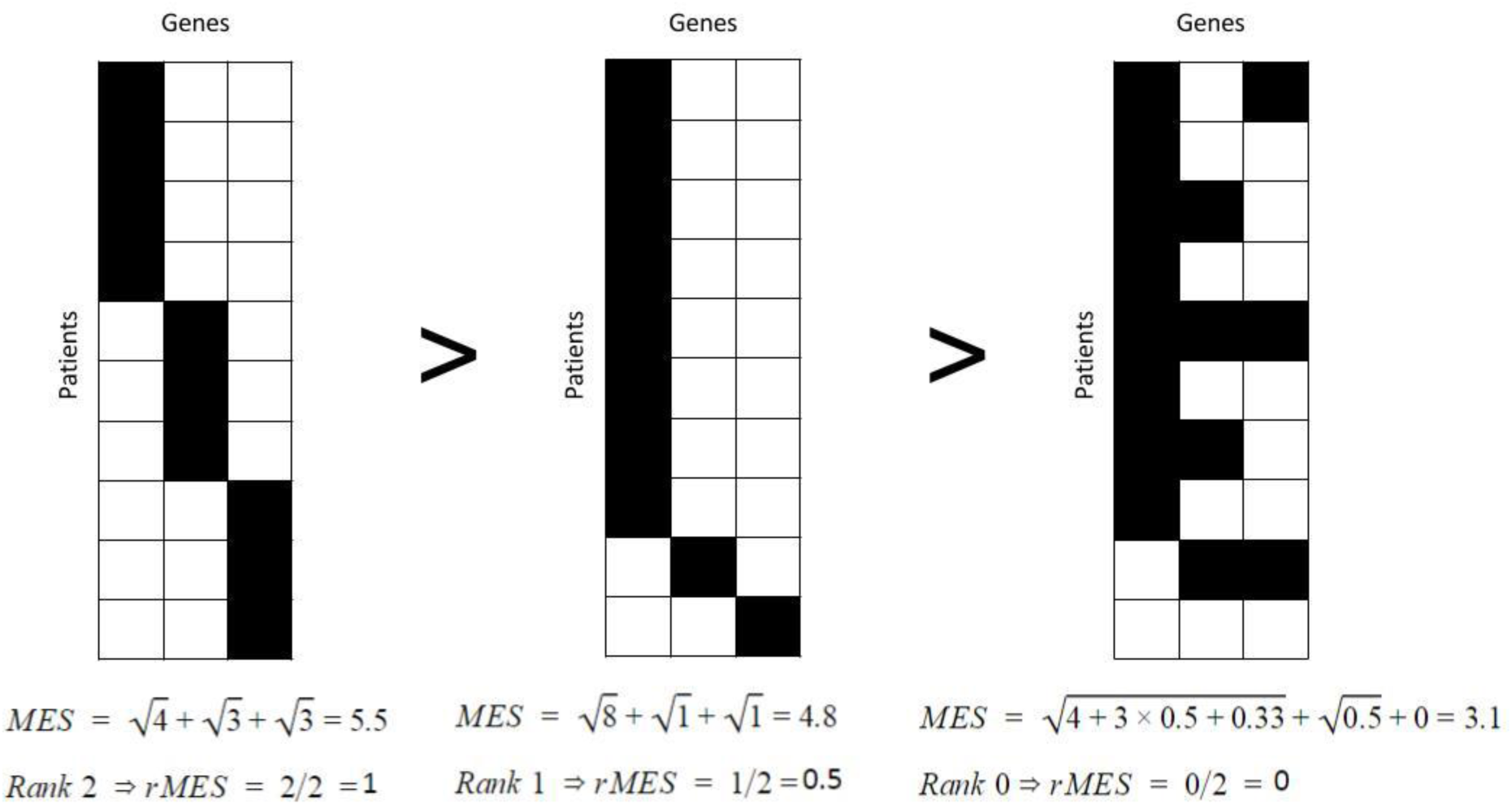
Calculation of *MES* and corresponding *rMES* scores for three different small subnetworks. Genes which make up the small subnetwork are represented as columns of matrices, patients are represented as rows. Genes with alterations in a specific patient are depicted as black tiles. Small subnetworks exhibiting perfect mutually exclusivity patterns (two most left small subnetworks) have higher *rMES* scores than small subnetworks with non-perfect mutual exclusivity patterns (most right small subnetwork). Also, small subnetworks having a more uniform distribution of gene alterations across patients have higher *rMES* scores as shown by the two most left small subnetworks.

#### Reinforced learning

Using the *rMES* for each small subnetwork, the reinforced learning step updates gene scores based on two parameters: *reinforcement* and *forgetfulness*. The *reinforcement* is a parameter that determines the maximal value by which a gene score can be increased in the next iteration. The reinforcement is multiplied by the highest *rMES* score of all small subnetworks to which the gene belongs, so the gene score of genes which are consistently in small subnetworks with high *rMES* scores will further increase with iterations.

The *forgetfulness* determines the fraction of the gene score that is retained in every subsequent iteration. This means that part of the gene score is effectively lost every iteration step and thus the gene scores of genes having persistently low *rMES* scores will go to zero. To calculate gene scores the following formula is used:

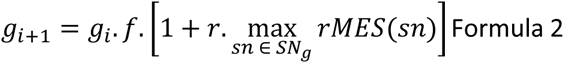

Where *g_i_* is the gene score at iteration *i*, *f* is the *forgetfulness, r* the *reinforcement, SN_g_* the set of small subnetworks containing the gene. If the gene score resulting from the formula is larger than 1, it is topped off at 1 as the maximal gene score can never be larger than 1. The default parameters of the method are forgetfulness *f* = 0.995, reinforcement *r =* 0.005 and 5000 iterations. In general, the sum of forgetfulness and reinforcement should be close to 1 and the reinforcement should be small (smaller than 0.01). This because small values for forgetfulness or large values for reinforcement would make the algorithm prone to stochastic effects. Note that genes which are not part of any small subnetwork are assigned a value of zero for 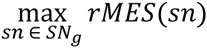.

In a final step we assign a rank to each gene that reflects to what extent a gene belongs to a mutual exclusivity pattern. Hereto we exploit the fact that genes belonging to a mutual exclusivity pattern tend to have a consistent increase in their gene score between iterations over time. Genes are ranked according to the maximal gene score they reach and in case of ties are based on how fast their score converges.

## Simulated data

To assess the performance of SSA.ME we used simulated data. The set of true positive driver genes was defined first by creating a target mutually exclusive pattern which in biological terms corresponds to a driver pathway. The target mutual exclusivity pattern was generated using a random walker with restart (5% restart chance) to select genes from the local network neighborhood of a randomly selected gene until 20 interactions have been visited in a high quality human reference network. This high quality human reference network was composed of HINT^27^, Interactome (HI-II-14)^28^ and Reactome^29^ interaction data.

To mimic real tumor data, we counted the number of mutated genes present in each tumor sample in the TCGA 2012 study and assigned an equal number of alterations to random genes, thus conserving the distribution of mutated genes per sample. We added mutually exclusive mutations to genes present in the target pattern in 30 % of the samples. Each sample had 5% chance to also be mutated in any of the other genes belonging to the same mutual exclusivity pattern as we allowed for “soft” mutual exclusivity patterns which are non-perfect across samples.

To evaluate the robustness of the method with respect to the used reference network, multiple simulated datasets were analyzed for different degrees of connectedness in the high quality human reference network: highly underconnected (50% of the edges were deleted from the reference network), mildly underconnected (25% of the edges deleted), lowly underconnected (10% edges deleted), true network (i.e. the high quality human reference network), lowly overconnected (10% additional random edges added to the reference network), mildly overconnected (25% additional edges) and highly overconnected (50% additional edges). We generated 100 different simulated datasets per network and ran SSA.ME. Performance was measured by receiver operating characteristic (ROC) curves.

To assess parameter sensitivity we tested the effect of using different parameter combinations on the performance. This included 400 simulations for all combinations of reinforcement *r* (from 0.0005 to 0.0100 in steps of 0.0005) and forgetfulness *f* (from 0.99 to 0.9995 in steps of 0.0005). Performance for each parameter combination was measured using the area under the curve (AUC).

## Breast Cancer TCGA Data

The TCGA Breast Cancer (BRCA) data published in 2012^2^ was downloaded from https://tcga-data.nci.nih.gov/docs/publications/brca_2012/. Level 2 files were used containing somatic mutations, RNA expression and copy number variations. Copy number alterations obtained from the original TCGA Breast cancer data were inferred with GISTIC^30^. In our analysis only genes in samples with high-level thresholds for amplifications/deletions and for which copy number alteration showed positive correlation with expression level were used. Priorization results were obtained by running SSA.ME on a non-stringently filtered input set, consisting of all genes having at least one genetic alteration (mutation or amplification/deletion correlated with expression) in the dataset. As a high quality human reference network we compiled information data from HINT^27^, Interactome (HI-II-14)^28^ and Reactome^29^.

Using the TCGA breast cancer data also allowed us to compare our results with the ones originally published by MEMo, a representative state-of-the-art method that searches for mutual exclusivity patterns using a reference network. To maximize the comparison, we ran our approach with the same reference network and with the same data as originally used by MEMo. This reference network is a non-curated reference network consisting of Reactome^29^, Panther^31^, KEGG^32^, INOH^33^ and interactions from non-curated sources (like high-throughput derived protein–protein interactions, gene co-expression, protein domain interaction, GO annotations, and text-mined protein interactions)^34^. Data were reproduced according to the description in the original paper, i.e. only retaining genes that were altered in at least ten samples. To illustrate how prioritizations were not largely affected by omitting this stringent prefiltering we redid the analysis in the same setting but using the less filtered input described above.

## Acknowledgements

The authors would like to thank Lieven Verbeke for the useful comments and discussions.

Funding: Ghent University Multidisciplinary Research Partnership ‘Bioinformatics: from nucleotides to networks’; Fonds Wetenschappelijk Onderzoek-Vlaanderen (FWO) [G.0329.09, 3G042813, G.0A53.15N]; Agentschap voor Innovatie door Wetenschap en Technologie (IWT) [NEMOA and the personal fellowship of Dries de Maeyer]; Katholieke Universiteit Leuven [PF/10/010] (NATAR)

## Additional Information

The author(s) declare no competing financial interests.

## Availability of materials and data

SSA.ME software is available at https://github.com/spulido99/SSA. The breast cancer dataset is available at its website https://tcga-data.nci.nih.gov/docs/publications/brca2012/. CADD scores version 1.3 were downloaded from http://cadd.gs.washington.edu/. The pan-cancer dataset is publicly available at https://www.synapse.org/#!Synapse:syn1710680.

## Author Contributions Statement

S.P.T, B.W, D.DM and K.M conceived the study. S.P.T, B.W. and D.DM developed the SSA framework. S.P.T and B.W developed, tested and analyzed the performance of the SSA application to mutual exclusivity. S.P.T, B.W and K.M wrote the manuscript. All authors reviewed the manuscript.

